# Common schizophrenia alleles are enriched in mutation-intolerant genes and maintained by background selection

**DOI:** 10.1101/068593

**Authors:** Antonio F. Pardiñas, Peter Holmans, Andrew J. Pocklington, Valentina Escott-Price, Stephan Ripke, Noa Carrera, Sophie E. Legge, Sophie Bishop, Darren Cameron, Marian L. Hamshere, Jun Han, Leon Hubbard, Amy Lynham, Kiran Mantripragada, Elliott Rees, James H. MacCabe, Steven A. McCarroll, Bernhard T. Baune, Gerome Breen, Enda M. Byrne, Udo Dannlowski, Thalia C. Eley, Caroline Hayward, Nicholas G. Martin, Andrew M. McIntosh, Robert Plomin, David J. Porteous, Naomi R. Wray, GERAD Consortium, David A. Collier, Dan Rujescu, George Kirov, Michael J. Owen, Michael C. O’Donovan, James T. R. Walters

## Abstract

Schizophrenia is a debilitating psychiatric condition often associated with poor quality of life and decreased life expectancy. Lack of progress in improving treatment outcomes has been attributed to limited knowledge of the underlying biology, although large-scale genomic studies have begun to provide such insight. We report the largest single cohort genome-wide association study of schizophrenia (11,260 cases and 24,542 controls) and through meta-analysis with existing data we identify 50 novel GWAS loci. Using gene-wide association statistics we implicate an additional set of 22 novel associations that map onto a single gene. We show for the first time that the common variant association signal is highly enriched among genes that are intolerant to loss of function mutations and that variants in these genes persist in the population despite the low fecundity associated with the disorder through the process of background selection. Associations point to novel areas of biology (e.g. metabotropic GABA-B signalling and acetyl cholinesterase), reinforce those implicated in earlier GWAS studies (e.g. calcium channel function), converge with earlier rare variants studies (e.g. *NRXN1*, GABAergic signalling), identify novel overlaps with autism (e.g. *RBFOX1*, *FOXP1*, *FOXG1*), and support early controversial candidate gene hypotheses (e.g. *ERBB4* implicating neuregulin signalling). We also demonstrate the involvement of six independent central nervous system functional gene sets in schizophrenia pathophysiology. These findings provide novel insights into the biology and genetic architecture of schizophrenia, highlight the importance of mutation intolerant genes and suggest a mechanism by which common risk variants are maintained in the population.

---

Schizophrenia is characterised by psychosis and negative symptoms such as social and emotional withdrawal. While onset of psychosis typically does not occur until late adolescence or early adult life, there is strong evidence from clinical and epidemiological studies that schizophrenia reflects a disturbance of neurodevelopment^1^. It confers substantial mortality and morbidity, with a mean reduction in life expectancy of 15-30 years^2,3^. Although recovery is possible, most patients have poor social and functional outcomes^4^. No substantial improvements in outcomes have emerged since the advent of antipsychotic medication in the mid-20th century, a fact that has been attributed to a lack of knowledge of pathophysiology^1^.

Schizophrenia is both highly heritable and polygenic, with risk ascribed to variants spanning the full spectrum of population frequencies^5-7^. The relative contributions of alleles of various frequencies is not fully resolved, but recent studies estimate that common alleles, captured by genome-wide association study (GWAS) arrays, capture between a third and a half of the genetic variance in liability^8^. There has been a long-standing debate, from an evolutionary standpoint, as to how common risk alleles might be maintained in the population, particularly given the early mortality and decreased fecundity associated with schizophrenia^9^. Various hypotheses have been proposed including compensatory advantage (balancing selection), whereby schizophrenia alleles confer reproductive advantages in particular contexts^10,11^; hitchhiking, whereby risk alleles are maintained by their linkage to positively selected alleles^12^; or contrasting theories that attribute these effects to rare variants and gene-environment interaction^13^. Addressing these competing hypotheses is now tractable given advances from recent studies of common genetic variation in schizophrenia.

The largest published schizophrenia GWAS, that from the Schizophrenia Working Group of the Psychiatric Genomics Consortium (PGC), identified 108 genome-wide significant loci and unequivocally demonstrated the value of increasing sample sizes for discovery in schizophrenia GWAS^5^. Here, we report by far the largest ancestrally and phenotypically homogeneous GWAS study of schizophrenia to date (N=11 260 cases, 24,542 controls). We combine these new data with the previous published GWAS to identify novel facets of genetic architecture and biology, and demonstrate that the evolutionary process of background selection can contribute to the maintenance of risk alleles in the population.

## RESULTS

### GWAS and Meta-analysis

We obtained genome-wide genotype information for the largest single-country study of schizophrenia (CLOZUK) based on 15,000 cases from the UK, 96% of whom were taking clozapine, an antipsychotic for treatment-resistant schizophrenia. Control datasets with UK ancestry were obtained from public repositories or through collaboration. To maximise homogeneity beyond that conferred by our ascertainment strategy, we set stringent thresholds for inclusion based on genotype-derived ancestry, resulting in a final CLOZUK GWAS of 11,260 cases and 24,542 controls (5,220 cases and 18,823 controls not in previous schizophrenia GWAS **- Methods; Figure S1; Figure S2**).

The CLOZUK GWAS analysis (λ_GC_=1.281, λ_1000_=1.018) yielded 18 genome-wide significant (GWS=p<5× 10^−8^) independent loci, four of which are novel (**Extended Data Table 1; Figure S3; Figure S4**). The CLOZUK SNP-based heritability (h^2^_SNP_) on the liability scale was 0.29 (assuming 1% prevalence; see **Extended Data Table 2** for a range of plausible prevalences) using linkage-disequilibrium score regression (LDSR)^14^. We demonstrate common genetic architecture between the CLOZUK and the independent PGC sample (excluding all CLOZUK samples) by showing high genetic correlation (95%), sign test enrichment (99% of PGC GWS SNPs show the same direction of effect in CLOZUK: p=2.04×10^−21^) and strong polygenic risk score overlap (p<1×10^−300^) – see **Methods** and Extended Data Table 3 and 4.

Having demonstrated high genetic correlation between datasets, we performed meta-analysis of the CLOZUK and the independent PGC dataset, excluding related and overlapping samples (total 40,675 cases and 64,643 controls, λ_GC_=1.586, λ_1000_=1.012; **Figure S5**). We identified 177 independently associated SNPs (**Extended Data Table 5**) that map to 143 independent loci (**Figure 1, Extended Data Table 6, Methods**). Overall, the association signal at the majority of previously identified loci strengthened in the meta-analysis (**Figure S6**), although 14 of the PGC loci were no longer genome-wide significant (**Extended Data Table 7**). Of the 52 loci not identified by the PGC, two have been reported as genome-wide significant in other studies: *ZEB2*^15^ and a locus on chromosome 8 (38.0-38.3 MB)^16^. Thus we identify 50 novel genetic loci for schizophrenia (**Extended Data Tables 6 and 8**).

**Figure 1.**
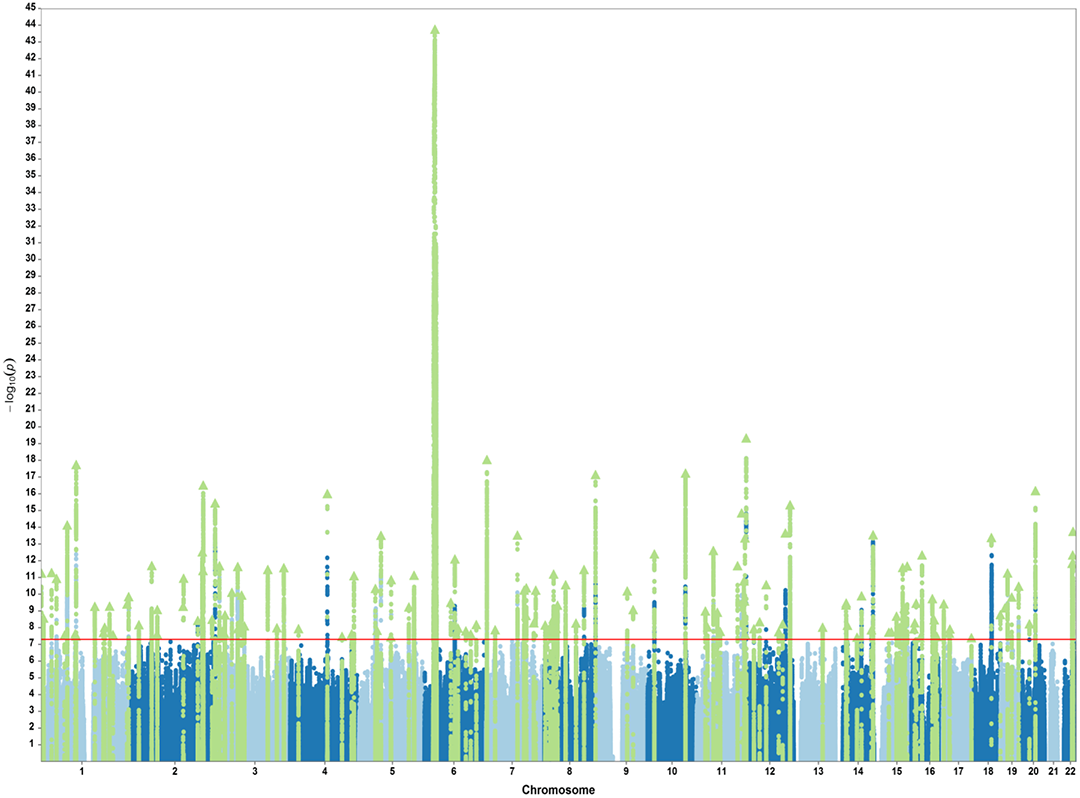
Manhattan plot of schizophrenia GWAS associations from the meta-analysis of CLOZUK and an independent PGC dataset (N=105318; 40675 cases and 64643 controls).

### Gene-based analysis

To exploit the additional information provided by multiple sub-genome-wide significant SNP associations within single genes, we undertook gene-wide analyses using MAGMA^17^ (**Methods**). In the full meta-analytic dataset, excluding the extended Major Histocompatibility region (xMHC) due to its complex LD structure, we identified 535 genome-wide significant genes (p<2.5×10^−6^), 240 of which did not co-localize with a genome-wide significant SNP locus from either the present meta-analysis or the PGC study^5^ (**Extended Data Table 9**). We then used the full genome-wide gene-based results to (i) investigate genetic architecture (ii) highlight single gene loci and (iii) conduct hypothesis and data-driven systems genomic analyses in schizophrenia.

### Mutation intolerant genes

Recent studies have shown that mutation intolerant genes capture much of the rare variant architecture of neurodevelopmental disorders such as autism, intellectual disability and developmental delay^18-20^. Work in parallel to the present study has shown that this extends to rare variants in schizophrenia (Singh et al, submitted). To investigate whether this holds for common variation, we analysed the set of loss-of-function (LoF) intolerant genes as defined by the Exome Aggregation Consortium (ExAC)^21^, using their preferred gene-level constraint metric, pLI ≥ 0.9 (probability of being LoF intolerant). Using gene set analysis in MAGMA, we found LoF intolerant genes (N=3230) were highly significantly enriched for schizophrenia common variant associations in comparison with all other genes (p=4.1×10^−16^).

To quantify the contribution of SNPs within LoF intolerant genes to schizophrenia SNP-based heritability (h^2^_SNP_) we conducted heritability analyses using partitioned LDSR^22^ (**Extended Data Table 10**). Overall, genic SNPs account for 64% of h^2^_SNP_, a 1.23-fold enrichment (compared to that expected from SNP content) whereas non-genic SNPs were depleted for SNP heritability content (h^2^_SNP_=33%; enrichment=0.69). Consistent with the analysis using MAGMA, h^2^_SNP_ was highly significantly enriched in LoF intolerant genes (2.01-fold; p=2.78×10^−24^). Common variation in LoF intolerant genes explained 30% of all h^2^_SNP_ (equating to 47% of all genic h^2^_SNP_) despite containing only 15% of all SNP content (29% of the total genic SNP content). In contrast, genes classed as non-LoF intolerant (pLI<0.9) were significantly depleted for h^2^_SNP_ relative to their SNP content (0.90-fold; p=5.86×10^−3^), although in absolute terms, SNPs in these genes accounted for 34% of h^2^_SNP_. A finer scale analysis of the relationship between LoF intolerance scores and enrichment for association showed that enrichment is restricted to genes with a pLI score above 0.9 (**Figure S7**).

### Common risk alleles maintained by background selection

Our novel finding that LoF intolerant genes, which by definition are under strong selective pressure, are enriched for common schizophrenia risk variants raises the question of how such alleles are maintained in the population. While the contribution of ultra rare variation in mutation intolerant genes to disorders associated with low fecundity can be accounted for by *de novo* mutation^23,24^, this cannot explain the persistence of common alleles. To address this question, we used partitioned LDSR to test the relationship between schizophrenia associated alleles and SNP-based signatures of natural selection. These included measures of positive selection, background selection, and Neanderthal introgression. We examined the heritability of SNPs after thresholding them at extreme values for these metrics (top 2%, 1% and 0.5%), while adjusting for baseline annotation sets including LoF intolerant genes (**Methods**).

We observed strong evidence for schizophrenia h^2^_SNP_ enrichment in SNPs under background selection (BGS), which was consistent across all the thresholds we examined (**Table 1**). We also found a significant and consistent depletion of h^2^_SNP_ in SNPs subject to positive selection as indexed by the CLR statistic. These two results are mutually consistent, as the calculation of the CLR statistic explicitly controls for B-statistic values^25^. This suggests that SNPs under positive selection, but under weak or no BGS, are depleted for association with schizophrenia. No significant relationship between h^2^_SNP_ and other positive selection or Neanderthal introgression measures was found after correction for multiple testing (**Table 1**).

**Table 1.** Heritability analysis of natural selection metrics. Partitioned LDSR regression results for SNPs thresholded by extreme values (defined as top percentiles vs all other SNPs) of each natural selection metric. All tests have been adjusted for 56 “baseline” annotations which include categories such as “LoF-intolerant” and “conserved” (see **Methods**). Negative enrichments should be considered zero (no contribution to h^2^_SNP_ by these SNPs). Underlined values indicate results surviving correction after adjusting for all tests (Bonferroni α=0.05/18=0.0028). Reference numbers for each metric indicate references in the main text.

|  |  | Top 2% of scores<br>(genome-wide) |  | Top 1% of scores<br>(genome-wide) |  | Top 0.5% of scores<br>(genome-wide) |  |
| --- | --- | --- | --- | --- | --- | --- | --- |
| Metric | Ref. | Enrichment | 2-sided<br>p-value | Enrichment | 2-sided<br>p-value | Enrichment | 2-sided<br>p-value |
| Background Selection (B-statistic) | 26 | <u>2.0612</u> | <u>0.0008</u> | <u>2.6477</u> | <u>1.2721x10<sup>-4</sup></u> | <u>2.8073</u> | <u>1.0226x10<sup>-4</sup></u> |
| Positive selection (CLR) | 25 | 0.6069 | 0.0506 | <u>0.3154</u> | <u>1.1349x10<sup>-5</sup></u> | <u>0.2382</u> | <u>1.1120x10<sup>-4</sup></u> |
| Positive selection (CMS) | 106 | 0.1703 | 0.0036 | 0.1902 | 0.0489 | -0.0108 | 0.0736 |
| Positive selection (XP-EEH) | 105 | 0.9918 | 0.9824 | 0.6211 | 0.4882 | -0.0018 | 0.1666 |
| Positive selection (iHS) | 104 | 0.8593 | 0.7095 | 0.7649 | 0.6992 | 1.3686 | 0.7130 |
| Neanderthal posterior probability (LA) | 107 | 0.7465 | 0.2175 | 0.7895 | 0.4930 | 0.9067 | 0.8428 |

To explicitly test the association of SNPs under background selection in LoF intolerant genes, we binned the B-statistic into four categories of increasing score, assigned SNPs in these bins to a LoF intolerant set, an “all other” genes set or a non-genic set (**Figure S8**). The lower boundary of the top bin (B-statistic >0.75) corresponds approximately to the top 2% BGS threshold in **Table 1** and is equivalent to a reduction in effective population size estimated at each SNP of 75% or more^26^. We found significant heritability enrichment across all B-statistic intervals in LoF intolerant genes which increased progressively with higher B-statistic scores. We also found enrichment for SNPs under BGS pressure in genes that are not LoF intolerant, although this was restricted to the highest B-statistic bin. There was no notable enrichment in any B-statistic intervals for non-genic regions.

### Notable single gene associations

In considering the novel GWAS loci and gene-based results from the meta-analysis we highlight the subset of the findings most likely pointing to single genes, given the difficulties in accurately locating causal genes for complex disorders in a GWAS framework^27^. These single gene associations were defined as GWAS SNP loci that span only a single gene +/-20 kb (N=22, 44% of novel loci, **Extended Data Table 6**) or MAGMA associated genes that were at least 100 kb from any other genome-wide significant gene (N=76, **Extended Data Table 9**). We acknowledge this does not imply that these are inevitably the pathogenic genes but their likelihood of being so is higher than that for genes in multigenic loci.

Details of the findings we consider most noteworthy are provided in **Table 2.** These results provide novel potential insights into schizophrenia pathogenesis as well as supporting existing hypotheses. To our knowledge we provide the first genome-wide significant findings supporting GABA involvement in schizophrenia pathogenesis through both SNP-based GWAS (*GABBR2*) and gene-based analysis (*SLC6A11*). Previous association studies have implicated calcium and cholinergic signalling involvement in schizophrenia^5,8^; we extend these findings by demonstrating association with an L-type calcium channel gene (*CACNA1D*) and with the acetyl cholinesterase gene (*ACHE*), a novel potential therapeutic target in schizophrenia which has also recently been implicated in autism through de novo mutation analysis^28^. We provide the first common variant support for *NRXN1*, previously implicated by single-gene copy number variation (CNV) in schizophrenia, autism and intellectual disability^28,29^ and we also highlight the involvement of other genes that increase risk across neurodevelopmental disorders including *RBFOX1*. Finally we find evidence for association for two genes whose functions are intimately linked to two of the most prominent genes from schizophrenia candidate gene era, *PDE4B*, a *DISC1* interactor^30^, and *ERBB4* a binding partner of neuregulin1 (*NRG1*)^31^.

**Table 2.**
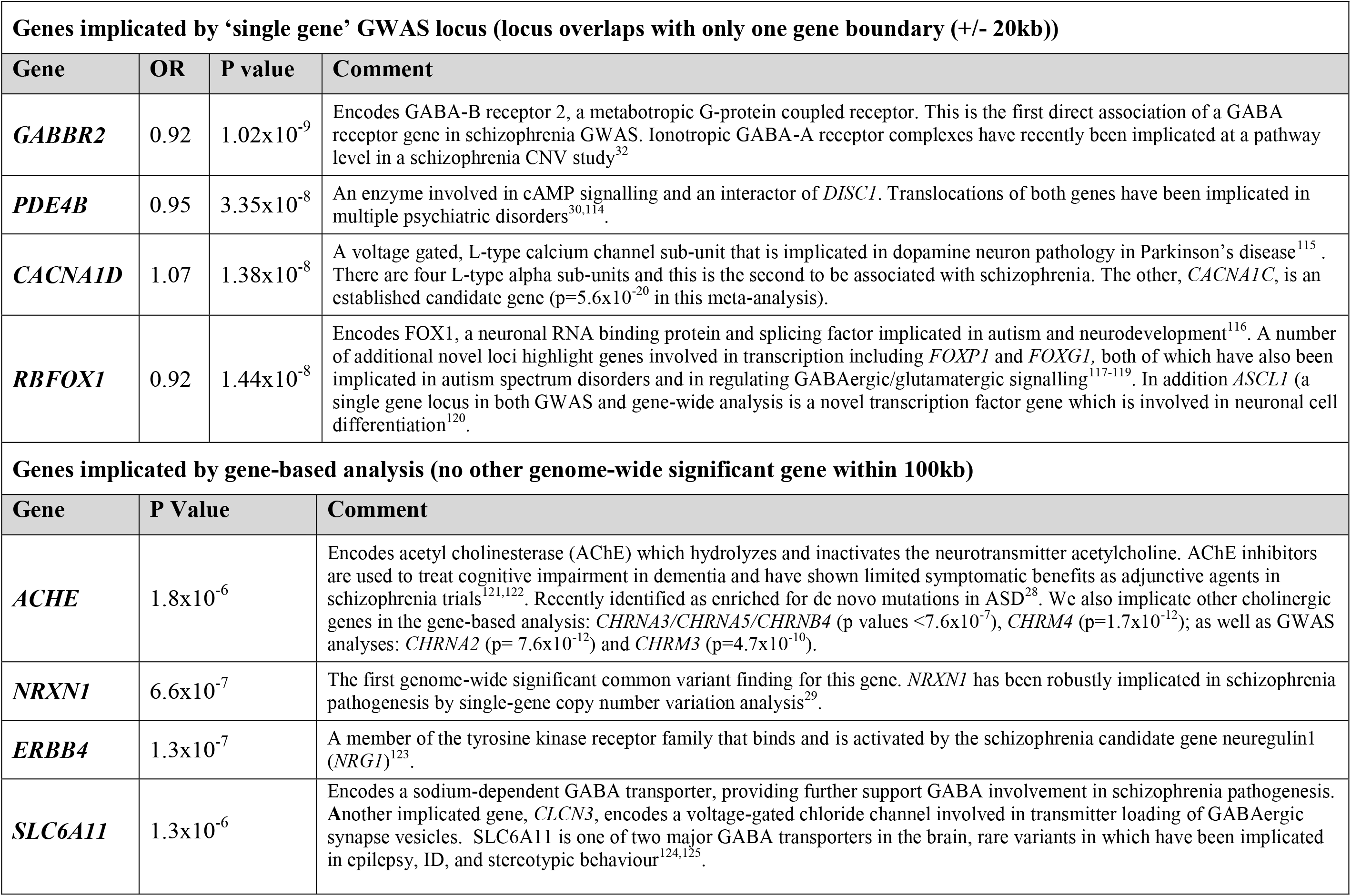
Genes with pre-existing genetic and functional hypotheses in schizophrenia or related neurodevelopmental disorders. Additional details an **Extended Data** Table 6 and **Extended Data** Table 9.

### Systems genomics

We undertook gene set analysis in MAGMA to gain insights into the biological systems underpinning schizophrenia. To maximise power we restricted our primary analysis to 134 gene sets we have previously postulated to be of likely relevance to the disorder on the basis of their involvement in the central nervous system, and which we subsequently showed effectively capture the excess CNV burden in schizophrenia^32^ (**Extended Data Table 11**). The set span subcellular neuronal function; neuronal cell physiology; cellular, brain region and fiber tract morphology; learning; behavior; and brain development, ^32^. In a GWAS context, we now show that collectively, this set captures a disproportionately high proportion of h^2^_SNP_ (30% of total heritability; enrichment=1.63; 46% of genic heritability; **Extended Data Table 10**).

Of the 134 sets, 54 were nominally significant of which 12 survived multiple-testing correction (family-wise error rate (FWER) p-value<0.05, **Extended Data Table 11**). Stepwise conditional analysis, adjusting sequentially for the more strongly associated gene sets, resulted in six gene sets that were independently associated with schizophrenia (**Table 3** and **Data Supplement**). These gene sets extend from low-level molecular and sub-cellular processes to broad behavioural phenotypes. The most strongly associated gene set is constituted by the targets of the Fragile X Mental Retardation Protein (FMRP)^33^. FMRP is a neuronal RNA-binding protein that interacts with polyribosomal mRNAs (the 842 target transcripts of this gene set^33^) and is thought to act by inhibiting translation of target mRNAs, including many transcripts of pre- and post-synaptic proteins. The FMRP target set has been shown to be enriched for rare mutational burden in *de novo* exome sequencing studies of autism^34^ and intellectual disability^35^. In schizophrenia studies, it has also been shown to be nominally significantly enriched for association signal in sequencing studies^8,35^ and in GWAS^5,8^ but only inconsistently in studies of copy number variation^32,36^. However, in none of these studies was enrichment at levels of significance that would be equivalent to genome wide significance (i.e. survive correction for a comprehensive testing of gene-sets in public databases). Thus we provide the most robust evidence to date for the involvement of this gene set in schizophrenia.

**Table 3.** Functional gene set analysis highlights six independent gene sets associated with schizophrenia FMRP: Fragile X Mental Retardation Protein. MP refers to Mammalian Phenotype Ontology term of the MGI: Mouse Genome Informatics (http://www.informatics.jax.org)37 from which gene sets were derived. FWER: Westfall-Young family-wise error rate as implemented in MAGMA^1787^. Conditional p value refers to stepwise conditional analysis that adjusts sequentially for ‘stronger’ associated gene sets.

| Gene Set | Number of genes | Enrichment p value (FWER) | Conditional p value |
| --- | --- | --- | --- |
| Targets of FMRP<sup>33</sup> | 798 | $1 \times 10^{-5}$ | $1.9 \times 10^{-8}$ |
| *Abnormal Behaviour*<br>(MP:0004924) | 1939 | $1.8 \times 10^{-4}$ | $1.4 \times 10^{-5}$ |
| 5-HT<sub>2C</sub> receptor complex<sup>39</sup> | 16 | 0.029 | 0.001 |
| *Abnormal Nervous System Electrophysiology*<br>(MP:0002272) | 201 | 0.003 | 0.002 |
| Voltage-gated calcium channel complexes<sup>38</sup> | 196 | 0.011 | 0.016 |
| *Abnormal Long Term Potentiation* (MP:0002207) | 142 | 0.030 | 0.031 |

We highlight another five gene sets that are independently associated with schizophrenia. Three of these derive from the Mouse Genome Informatics database^37^ and relate to behavioural and neurophysiological correlates of learning; Abnormal Behaviour (MP:0004924), Abnormal Nervous System Electrophysiology (MP:0002272) and Abnormal Long Term Potentiation (MP:0002207). We note that two of these gene sets (MP:0004924 and MP:0002207) were among the five most enriched of 134 gene sets tested in a recent schizophrenia CNV analysis^32^. The remaining two independently associated genes sets were voltage-gated calcium channel complexes^38^ and the 5-HT_2C_ receptor complex^39^. The calcium channel finding confirms extensive evidence from common and rare variant studies implicating calcium channel genes in schizophrenia^5,8^, including a novel GWAS locus in *CACNA1D* identified in our meta-analysis. Whilst there is less convergent evidence in support of the involvement of the 5-HT_2C_ receptor complex in schizophrenia, the fact that we identify independent association for this gene set implicates these genes in schizophrenia pathophysiolology and potentially rejuvenates a previous avenue of 5-HT_2C_ ligand therapeutic endeavour in schizophrenia research^40^. However we interpret this result with caution given the small size of this gene set and the fact that a number of its genes encode synaptic proteins that are structurally related to other receptor complexes^39^, not only 5-HT_2C_.

### Systems genomics and mutation intolerant genes

Together, LoF intolerant genes and the conditionally independent (“significant”) CNS-related gene sets together account for 39% of schizophrenia SNP-based heritability, equating to 61% of genic heritability (**Figure 2A**; Extended Data Table 10). This is likely to be an underestimation of the true effect of these gene sets since distal non-genic regulatory elements (not included in this analysis) will add to the heritability explained by these genes. In examining the relationship between the two gene sets (**Figure 2A**), genes belonging to both categories were the most highly enriched (2.6-fold; p=7.90×10^−15^), although LoF intolerant genes that were not annotated to our significant CNS gene sets still displayed robust enrichment for SNP-based heritability (1.74-fold; p=9.77×10^−10^), while genes that were in the significant CNS gene sets but had pLI<0.9 showed more modest enrichment (1.39-fold; p=6.05×10^−4^). Notably genes outside these categories showed a depletion in heritability relative to their SNP content (enrichment=0.79, p=1.82×10^−7^).

**Figure 2.**
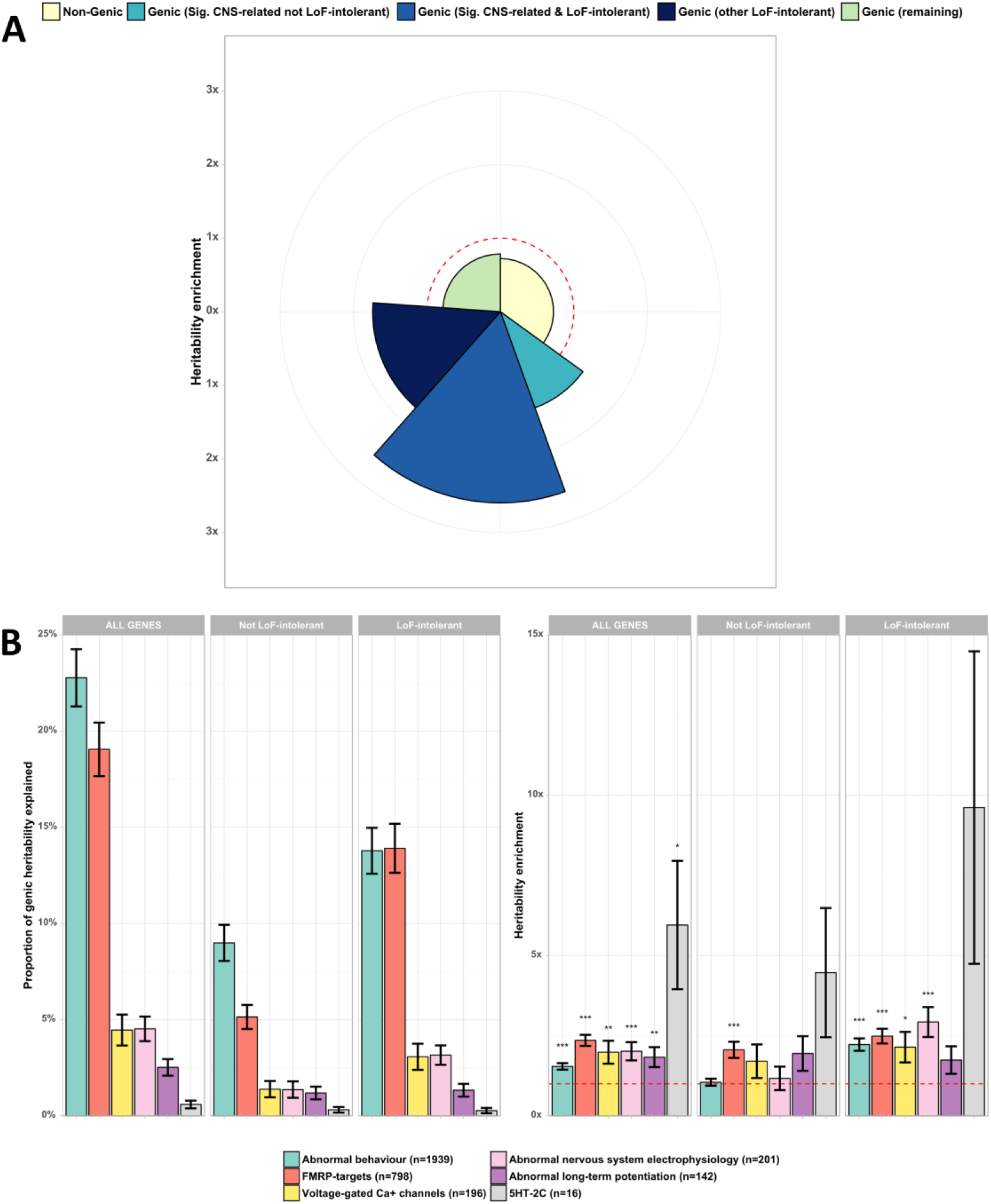
Partitioned heritability analysis of gene sets in schizophrenia. **A**: Heritability of genomic partitions and the six conditionally independent (“significant”) gene sets (**Table 3**). Radius of each segment indicates the degree of enrichment while the arc (angle of each slice) indicates the percentage of total SNP-based heritability explained. No relative enrichment or depletion (enrichment=1) is shown by the dashed red line. **B**: Heritability of the significant CNS gene sets dissected by their overlap with LoF-intolerant genes. Asterisks indicate the significance of each heritability enrichment (* <=0.05; ** <=0.01; *** <=0.001).

This general pattern remained when we focussed on the six significant CNS gene sets individually, in that the enrichment in these gene sets derives primarily from their intersection with LoF intolerant genes (**Figure** 2B). Indeed, only the targets of FMRP showed significant enrichment for SNPs in genes that are not LoF intolerant (2.06-fold; p=4.23×10^−5^).

### Data-driven gene set analysis

To set the systems genomics results in context, we undertook a purely data-driven analysis of a larger comprehensive annotation of gene sets from multiple public databases, totalling 6,677 gene sets (**Methods, Extended Data Table 12**). The LoF intolerant gene set was the most strongly enriched followed by the two strongest associated functional gene sets we had specified in our CNS gene set analysis (FMRP targets and MGI Abnormal Behaviour genes). The other three gene sets that survive FWER correction and conditional analyses are calcium ion import (GO:0070509), membrane depolarisation during action potential (GO:0086010) and synaptic transmission (GO:0007268). These sets show a marked functional and genic overlap with the independently associated sets from our primary CNS systems genomic analysis. Indeed if we repeat the data-driven comprehensive gene set analysis whilst adjusting for the six independently associated CNS gene sets, then the only surviving enrichment term is the LoF intolerant genes. These results are consistent with those from CNV analysis^32^ in that they do not support annotations other than those related to CNS function, and demonstrate that hypothesis based analysis to maximise power does not substantially impact on the overall pattern of results.

## DISCUSSION

In the largest genetic study of schizophrenia to date, we identify 50 novel genomic loci and a further 22 gene-wide associations that map onto to a single gene. We also explore the genomic architecture of, and the evolutionary pressures on, common variant associations to the disorder, and conduct extensive gene-based and systems genomic analyses. As discussed above and in table 1, novel findings point to GABAegic and cholinergic transmission as relevant to the aetiopathogenesis of schizophrenia. We widen the array of calcium channel receptors implicated in the disorder, and identify associations that support historical candidate gene networks that have been the subject of intensive functional investigation, but which have remained controversial in the absence of support from large scale genome-wide studies (*PDE4B*, *ERBB4*). We demonstrate congruence between common variant implicated genes and multiple rare variant associations to neurodevelopmental disorders, including *NRXN1*, *RBFOX1*, and *FOXG1* indicating that studies of the impact of rare mutations in these genes are likely to be relevant to schizophrenia pathogenesis more widely. Systems genomic analysis highlights six gene sets that are independently associated with schizophrenia, and point to molecular, physiological and behavioural pathways involved in schizophrenia pathogenesis. We also provide the first support at a genome-wide significant equivalent level implicating targets of FMRP in the disorder.

Our study provides the first evidence linking common variation in loss-of-function intolerant genes to risk of developing schizophrenia and demonstrates that these genes account for a substantial proportion (30%) of schizophrenia SNP-based heritability. Our findings underline the importance and value of well-curated catalogues of human genetic diversity and indicate that such resources have relevance for common variation and common disease. The Exome Aggregation Consortium^21^ reported that pLI score is positively correlated with gene expression across tissues, and this is also true for brain expression^41^. Furthermore high pLI genes have more protein-protein interaction partners, and pathways with the highest median pLI scores are ‘hub’ processes such as spliceosome and proteasome^21^. This evidence points to LoF intolerant genes being involved in core biological processes and presumably explains why they are under strong selective pressure. Given mutation intolerance and high selection pressure, our finding that common schizophrenia risk variants are enriched in loss-of-function intolerant genes appears paradoxical. However, we also present novel evidence that the major mechanism responsible for the persistence of common schizophrenia variation, both in LoF intolerant genes and across the genome, is background selection (BGS).

Selection against deleterious variants (purifying selection) is the main driver of genomic conservation over evolutionary timescales^42^. Regions of the genome under strong purifying selection constitute around 8% of the human genome^43^, and primarily include sequences which are important functionally and associated with fitness and survival^44^. BGS is a consequence of purifying selection occurring in regions of low recombination^45,46^. In such regions, selection against strongly deleterious variants causes whole haplotypes to be removed from the gene pool, which reduces genetic diversity at the locus in a manner equivalent to a reduction in effective population size^47^. This has the effect of impairing the efficiency of the selective process, allowing alleles with neutral or small deleterious effects that are not on haplotypes carrying strongly deleterious variants to rise in frequency by drift^45^. Our finding that schizophrenia-associated common SNPs are associated with high levels of BGS suggests a mechanism by which such variants, even if they are mildly deleterious, can be maintained in the population. The pruning of haplotypes carrying strongly deleterious mutations, which is expected to be particularly intense in LoF intolerant genes^21^, would allow these variants to persist in the population for as long as they reside on a haplotype free of these mutations. High LD additionally inhibits their amalgamation or replacement by recombination^45,48^. Such mechanisms of selection at linked sites have been shown to be compatible with the current knowledge about the genomic architecture of complex human traits^49^ and to influence phenotypes in model organisms, including gene expression^50^.

We did not find enrichment for any measure of positive selection or Neanderthal introgression. A recent study explained a negative correlation between schizophrenia associations and metrics indicative of a Neanderthal selective sweep as evidence for positive selection or polygenic adaptation in schizophrenia^12^. We do not replicate this correlation in our model, which addresses the contribution of BGS, and hence our results are not consistent with large contributions of positive selection to the genetic architecture of schizophrenia (**Table 1**). Indeed positive selection is not widespread in humans, as reported by other studies that explicitly considered or accounted for BGS^25,51^. Polygenic adaptation, the co-occurrence of many subtle allele frequency shifts at loci influencing complex traits^52^, though an intriguing possibility, has not been implicated in psychiatric phenotypes, including schizophrenia, in recent analyses^53,54^. In contrast, BGS has been proposed as a mechanism driving Human-Neanderthal incompatibilities, as regions with higher estimated B-statistics have lower estimated Neanderthal introgression^55^. We therefore conclude that the bulk of the BGS signal we obtain is unlikely to be influenced by positive selection^26^ and that previous results linking positive selection and genetic variation in schizophrenia are likely to have been confounded by failure to control for BGS^46^.

In summary, we demonstrate the capacity and continued importance of improving the power of genome-wide association studies for answering fundamental questions about the nature of schizophrenia. We identify large numbers of novel associations that both suggest novel, and support established, hypotheses about the biology of schizophrenia. We show convergence of risk variants on mutation intolerant genes, and identify background selection as the main mechanism by which common schizophrenia-related variation is maintained in the population.

## METHODS

### Case sample collection

We collected blood samples from those with treatment-resistant schizophrenia (TRS) in the UK through the mandatory clozapine blood-monitoring system for those taking clozapine, an antipsychotic licensed for TRS. Following national research ethics approval and in line with UK Human Tissue Act regulations we worked in partnership with the commercial companies that manufacture and monitor clozapine in the UK. We ascertained anonymous aliquots of the blood samples collected as part of the regular blood monitoring that takes place whilst taking clozapine due to a rare haematological adverse effect, agranulocytosis. The CLOZUK1 sample was assembled in collaboration with Novartis (Basel, Switzerland). The company, through their proprietary Clozaril® Patient Monitoring Service (CPMS), provided whole-blood samples and anonymised phenotypic information for 6,882 individuals with TRS (5528 cases post-QC), which were included in the in a recent schizophrenia GWAS by the PGC^5^. The CLOZUK2 sample, previously unreported, was assembled in collaboration with the other major company involved in the supply and monitoring of clozapine in the UK, Leyden Delta (Nijmegen, Netherlands). The company, through their proprietary Zaponex® Treatment Access System (ZTAS), provided whole-blood samples and anonymised phenotypic information for 7,417 of those taking clozapine (4973 cases post-QC). Both Clozaril® and Zaponex® are bioequivalent brands of clozapine licensed in the UK^56^.

We restricted the CLOZUK1 and CLOZUK2 samples to those with a clinician reported diagnosis of treatment-resistant schizophrenia. The UK National Institute for Health and Care Excellence (NICE) advise prescription of clozapine is reserved for those with schizophrenia in whom two trials of antipsychotics have failed (including one second-generation antipsychotic)^57^ which mirrors the criteria for licensed use of clozapine. The sole alternative licensed indication for clozapine in the UK is for the management of resistant psychosis in Parkinson’s disease (PD)^58^ and, although this is a rare indication, we excluded PD patients (n=8) from the case dataset. We also excluded those with off-license indications, which included those with alternative clinician diagnoses of bipolar affective disorder and personality disorders (n=56). Together with the clinical guidelines outlined, these exclusions ensure that CLOZUK1 and CLOZUK2 samples are from those patients that conform to a clinical description of TRS. We have reported the use of CLOZUK1 as a schizophrenia dataset in previous publications^5,7,59,60^ and have presented evidence to support the use of TRS-defined individuals as valid schizophrenia samples^32^, which we have updated and present in **Supplementary Note 1,** including validation of a clinician diagnosis of TRS against research diagnostic criteria for schizophrenia.

In addition we also included in our analysis a more conventional cohort of UK-based patients with schizophrenia (CardiffCOGS). Recruitment was via secondary care, mainly outpatient, NHS mental health services in Wales and England. These patients were not exclusively taking clozapine at the time of their recruitment. All cases underwent a SCAN interview^61^ and case note review followed by consensus research diagnostic procedures and were included if they had a DSM-IV schizophrenia or schizoaffective disorder-depressive type diagnosis, as previously reported^7^. The CardiffCOGS samples were recruited and genotyped in two waves: CardiffCOGS1 (512 cases, included in a previous GWAS^5^) and CardiffCOGS2 (247 cases).

Genotyping for these case samples was performed by the Broad Institute (Massachusetts, USA) for the CLOZUK1 sample and CardiffCOGS1 cases, using Illumina HumanOmniExpress-12 and OmniExpressExome-8 chips as described elsewhere^7^. The CardiffCOGS2 cases and the CLOZUK2 sample were genotyped by deCODE Genetics (Reykjavík, Iceland), using Illumina HumanOmniExpress-12 chips.

As all of these samples are intrinsically related and their recruitment and genotyping conforms to clinical and technical standards, thus we have combined them and used the term “CLOZUK” throughout this manuscript to describe the schizophrenia case dataset.

### Control sample collection

Control samples were collected from publicly available sources (EGA) or through collaboration with the holders of the datasets. Individual datasets were curated using the same procedures as the case-only datasets. In order to maximize the numbers of individuals that could be effectively included in the GWAS without introducing confounders, these datasets were chosen on the basis of having recruited individuals with self-reported UK ancestry (either exclusively or primarily) and having been genotyped on Illumina chips. A summarized view of all the datasets included in the GWAS is provided in **Supplementary Note 2,** which includes further details of the control datasets.

### Genotype quality-control (QC)

Given the many data sources used and the variety of genotyping chips available, a stringent quality control (allowing only 2% of missing SNP and individual data) was performed separately in each individual dataset, using PLINK v1.9^62^ and following standard procedures^63^. To facilitate merging and to avoid common sources of batch effects^64^, all SNPs in each dataset were also aligned to the plus strand of the human genome (build 37p13), removing strand-ambiguous markers in the process. As most control datasets lacked any markers in the X and Y chromosomes, or in the mitochondrial DNA, every SNP from these regions was discarded in the combined genotype data. The final merge of all case and control datasets left 203,436 overlapping autosomal SNPs.

All individuals were imputed simultaneously in the Cardiff University high-performance computing cluster RAVEN^65^, using the SHAPEIT/IMPUTE2 algorithms^66,67^. As reference panels, a combination of the 1000 Genomes phase 3 (1KGPp3) and UK10 K datasets was used, as this has previously been shown to increase the accuracy of imputation for individuals of British ancestry, particularly for rare variants^68^.

After imputation, a principal component analysis (PCA) of common variants (MAF higher than 5%) was carried out to obtain a general summary of the population structure of the sample, using the EIGENSOFT v6 toolset^69^. A plot of the first two PCs showed the existence of a large fraction of cases (∼20%) with no overlapping controls (**Figure S1, A**). A comparison with the 1KGPp3 dataset, performed using PCA and ADMIXTURE^70^ estimates, showed that most of these cases were similar in genetic ancestry to non-European individuals, namely from the East Asian or West African superpopulations (**Figure S1, B**). In order to ameliorate population stratification in the association analysis^71^, all individuals not falling into an area delimited by the mean and 3 standard deviations of the two first principal components of the control samples were excluded from further analyses (**Figure S1, C**). By repeating PCA only on the selected individuals, no outliers could be detected in the first two principal components, and ADMIXTURE plots were homogenised as well (**Figure S2**).

The CLOZUK sample was further pruned by removing all individuals with inbreeding coefficients (F) higher than 0.2, and leaving only a random member of each pair with a relatedness coefficient (π̂) higher than 0.2. Furthermore, to ensure the independence of our analyses with previous GWAS conducted by the Schizophrenia Working Group of the PGC, relatedness coefficients of CLOZUK individuals were also calculated with all the individual datasets included in the latest PGC GWAS^5^ following approval by the Consortium. Detected genetic relatives (or duplicates) were excluded in CLOZUK in the same way as intra-population relatives. After this imputation and curation process, 35,802 samples (11,260 cases and 24,542 controls) with 9.65 million imputed markers (INFO > 0.3 and MAF > 0.001) remained in the CLOZUK dataset.

### GWAS and reporting of independently-associated regions

The CLOZUK schizophrenia GWAS was performed using logistic regression with imputation probabilities (“dosages”) adjusted for 11 PCA covariates. These covariates were chosen as those nominally significant (p<0.05) in a logistic regression for association with the phenotype^72^. To avoid overburdening the GWAS power by adding too many covariates to the regression model^73^, only the first twenty PCs were considered and tested for inclusion, as higher numbers of PCs only become useful for the analysis of populations that bear strong signatures of complex admixture^74^. The final set of covariates included the first five PCs (as recommended for most GWAS approaches^75^) and PCs 6, 9, 11, 12, 13 and 19. Quantile-quantile (QQ) and Manhattan plots are shown in **Figure S3** and **S4**.

In order to identify independent signals among the regression results, signals were amalgamated into putative associated loci using the same two-step strategy and parameters as PGC (**Extended Data Table 1**). In this procedure, regular LD-clumping is performed (r^2^=0.1, p<1×10^−4^; window size <3 Mb) in order to obtain independent index-SNPs. Afterwards loci were defined for each index SNP as the genomic region which contains all other imputed SNPs within an r^2^≥0.6. To avoid inflating the number of signals in gene-dense regions or in those with complex LD, all loci within 250 kb of each other were annealed.

### Meta-analysis with PGC

A total of 6,040 cases and 5,719 controls from CLOZUK were included in the recent PGC study^5^. We reanalysed the PGC data after excluding all these cases and controls, obtaining a sample termed ‘INDEPENDENT PGC’ (29,415 cases and 40,101 controls). Adding the summary statistics from this independent sample to the CLOZUK GWAS results allowed for a combined analysis of 40,675 cases and 64,643 controls (without duplicates or related samples). This meta-analysis was performed using the fixed-effects procedure in METAL^76^ with weights derived from standard errors. For consistency with the PGC analysis, additional filters (INFO > 0.6 and MAF > 0.01) were applied to the CLOZUK and INDEPENDENT PGC summary statistics, leaving 8 million markers in the final meta-analysis results. QQ and Manhattan plots are shown in **Figure S5** and **Figure 2**. The same procedure as above was used in order to report independent loci from this analysis (**Extended Data Table 5**, **Extended Data Table 6**). As raw PGC genotypes were not available for the LD-clumping procedure, 1KGPp3 was used as a reference.

### Estimation and assessment of a polygenic signal

Association signals caused by the vast polygenicity underlying complex traits can be hard to distinguish from confounders related to sample relatedness and population stratification. In order to effectively disentangle this issue, we used the software LD-Score v1.0 to analyse the summary statistics of the CLOZUK GWAS, and estimate the contribution of confounding biases to our results by LDSR^14^. An LD-reference was generated using 1KGPp3 using the standard parameters implemented in the software. In order to improve accuracy, indels were discarded and markers used in this procedure were restricted to those with INFO > 0.9 and MAF > 0.01, a total of 5.16 million SNPs. The resulting LD-score intercept was 1.085 ± 0.010, which compared to a mean χ^2^ of 1.417 indicates a polygenic signal contribution of at least 80%. This is in line with other well-powered GWAS studies of complex human traits^14^, including schizophrenia^5^. SNP-based heritability (h^2^_SNP_) in our datasets (CLOZUK, INDEPENDENT PGC and the CLOZUK + PGC meta-analysis) was also calculated in this analysis, and transformed to a liability scale using a population prevalence of 1% (registry-based lifetime prevalence^77^). For reference and compatibility with epidemiological studies of schizophrenia, prevalence estimates of 0.7% (lifetime morbid risk^78^) and 0.4% (point prevalence^78^) were used for additional liability-scale h^2^_SNP_ calculations (**Extended Data Table 2**).

We also used LD-Score to compare the genetic architecture of CLOZUK and INDEPENDENT PGC, by calculating the correlation of their summary statistics^79^. A genetic correlation coefficient of 0.9541 ± 0.0297 was obtained, with a p-value of 6.63×10^−227^. There were 76 independent SNPs at a genome-wide significant (GWS) level in the INDEPENDENT PGC dataset after excluding the extended major histocompatibility complex region (xMHC). Using binomial sign tests based on clumped subsets of SNPs we found all but 1 (98.6%) of the 76 GWS SNPs from the INDEPENDENT PGC were associated with the same direction of effect in the CLOZUK sample, a result highly unlikely to reflect chance^80^ (p=2.04×10^−21^, **Extended Data Table 3**). Even of those with an association p-value less than 10^−4^ in the INDEPENDENT PGC sample, 82% showed enrichment in the CLOZUK cases (p=3.44×10^−113^), confirming very large numbers of true associations will be discovered amongst these SNPs with increased sample sizes. Additionally, the new sample introduced in this study (CLOZUK2) was compared by the same methods with the independent PGC dataset and showed results consistent with the full CLOZUK analysis, providing molecular validation of this sample as a schizophrenia sample (**Extended Data Table 3**).

We went on to conduct polygenic risk score analysis. Polygenic scores for CLOZUK were generated from INDEPENDENT PGC as a training set, using the same parameters for risk profile score (RPS) analysis in PGC^5^, arriving at a high-confidence set of SNPs for RPS estimation by removing the xMHC region, indels, and applying INFO > 0.9 and MAF > 0.1 cut-offs. Scores were generated from the imputation dosage data, using a range of p-value thresholds for SNP inclusion^81^ (5×10^−8^, 1×10^−5^, 0.001, 0.05 and 0.5). In this way, we can assess the presence of a progressively increasing signal-to-noise ratio in relation to the number of markers included^82^. As in the PGC study, we find the best p-value threshold for discrimination to be 0.05 and report highly significant polygenic overlap between the INDEPENDENT PGC and CLOZUK samples (p<1×10^−300^, r^2^=0.12, **Extended Data Table 4**), confirming the validity of combining the datasets. For comparison with other studies we also report polygenic variance on the liability scale^83^, which amounted to 5.7% for CLOZUK at the 0.05 p-value threshold (**Extended Data Table 4**). As in the PGC study the limited r^2^ and AUC in this analysis restricts the current clinical utility of these scores in schizophrenia.

### Gene set analysis

In order to assess the enrichment of sets of functionally related genes, we used MAGMA v1.03^84^ on the CLOZUK + PGC meta-analysis summary statistics, after excluding the xMHC region. First, gene-wide p-values were calculated by combining the p-values of all SNPs inside genes after accounting for linkage disequilibrium (LD) and outliers. This was performed allowing for a window of 35 kb upstream and 10 kb downstream of each gene in order to capture the signal of nearby SNPs that could fall in regulatory regions^85,86^. Next we calculated competitive gene set p-values on the gene-wide p-values after accounting for gene size, gene set density and LD between genes. For multiple testing correction in each gene set collection, a FWER^87^ was computed using 100,000 re-samplings.

We performed sequential analyses using three approaches:

1. **Loss-of-function intolerant genes:** We tested the enrichment of the loss-of-function (LoF) intolerant genes described by ExAC ^21^. This set comprises all genes defined in the ExAC database^88^ as having a probability of LoF-intolerance (pLI) statistic higher than 90%. While these genes do not form part of cohesive biological processes or phenotypes, they have been previously found to be highly expressed across tissues and developmental stages^21^. Also, they are enriched for hub proteins^89^, which makes them interesting candidates for involvement in the “evolutionary canalisation” processes that have been proposed to lead to pleiotropic, complex disorders^90^.
2. **CNS-related genes:** These gene sets were compiled in our recent study^32^, and include 134 gene sets related to different to aspects of central nervous system function and development. These include, among others, gene sets which have been implicated in schizophrenia by at least two independent large-scale sequencing studies^8,35^: targets of the fragile-X mental retardation protein (FMRP^33^), constituents of the N-methyl-D-aspartate receptor (NMDAR^91^) and activity-regulated cytoskeleton-associated protein complexes (ARC^92,93^), as well as CNS and behavioural gene sets from the Mouse Genome Informatics database version 6 (http://www.informatics.jax.org) ^37^.
3. **Data-driven:** The final systems genomic analysis was designed as an “agnostic” approach, and thus a large number of gene sets from different public sources was included, which summed to a total of 6,677 sets. Gene set sources were selected to encompass a comprehensive collection of biochemical pathways and gene regulatory interaction networks, not necessarily conceptually related to psychiatric disorders. Similar approaches have been successful elsewhere^86,94^. In building this analytic approach, the LoF intolerant gene set and all sets in the *CNS-related* collection were used. Additionally, 2,693 gene sets with direct experimental evidence and a size of 10-200 genes^86^ were extracted from the Gene Ontology (GO^95^) database release 01/02/2016; 1,787 gene sets were extracted from the 4^th^ ontology level of the Mouse Genome Informatics database version 6; 1,585 gene sets were extracted from REACTOME^96^ version 55; 290 gene sets were extracted from KEGG^97^ release 04/2015; and 187 gene sets were extracted from OMIM^98^ release 01/02/2016.

Detailed results of the analyses of the CNS-related and data-driven collection are given in **Extended Data Table 11** and **Extended Data Table 12**. Reported numbers of genes in each gene set are those with available data in the meta-analysis. This may differ from the original gene set description as some genic regions (such as those in the X chromosome) had null or poor SNP coverage. The gene content of the CNS-related gene sets that survive conditional analysis (“significant”) is given in MAGMA format in the **Data Supplement**.

### Partitioned heritability analysis of gene sets

It is known that the power of a gene set analysis is closely related to the total heritability of the phenotype and the specific heritability attributable to the tested gene set^99^. In order to assess the heritability explained by the genes carried forward after the main gene set analysis, LD-Score was again used to compute a partitioned heritability estimate of CLOZUK + PGC using the gene sets as SNP annotations. As in the MAGMA analysis, the xMHC region was excluded from the summary statistics. These were also trimmed to contain no indels, and only markers with INFO > 0.9 and MAF > 0.01, for a total of 4.64 million SNPs. As a recognised caveat of this procedure is that model misspecification can inflate the partitioned heritability estimates^22^, all gene sets were annotated twice: Once using their exact genomic coordinates (extracted from the NCBI RefSeq database^100^) and another with regulatory regions taken into account using the same upstream/downstream windows as in the MAGMA analyses. Additionally, all SNPs not directly covered by our gene sets of interest were explicitly included into other annotations (“non-genic”, “genic but not LoF-intolerant”) based on their genomic location. Finally, the “baseline” set of 53 annotations from Finucane et al. 2015^22^, which recapitulates important molecular properties such as presence of enhancers or phylogenetic conservation, was also incorporated in the model. All of these annotations were then tested jointly for heritability enrichment. We note that using exact genic coordinates or adding regulatory regions made little difference to the estimated enrichment of our gene sets, and thus throughout the manuscript we report the latter for consistency with the MAGMA gene set analysis (**Figure 2; Extended Data Table 10**).

### Natural selection analyses

We aimed to explore the hypothesis that some form of natural selection is linked to the maintenance of common genetic risk in schizophrenia^101-103^. In order to do this, for all SNPs included in the CLOZUK + PGC meta-analysis summary statistics, we obtained four different genome-wide metrics of positive selection (iHS^104^, XP-EEH^105^, CMS^106^ and CLR^25^), one of background selection (B-statistic^26^, post-processed by Huber et al. 2016^25^) and one of Neanderthal introgression (average posterior probability LA^107^). The use of different statistics is motivated by the fact that each of them is tailored to detect a particular selective process that acted on a particular timeframe (see Vitti et al. 2013^51^ for a review). For example, iHS and CMS are based on the inference of abnormally long haplotypes, and thus are better powered to detect recent selective sweeps that occurred during the last ∼30,000 years^108^, such as those linked to lactose tolerance or pathogen response^106^. On the other hand, CLR incorporates information about the spatial pattern of genomic variability (the site frequency spectrum^109^), and corrects explicitly for evidences of background selection, thus being able to detect signals from 60,000 to 240,000 years ago^25^. The B-statistic uses phylogenetic information from other primates (chimpanzee, gorilla, orang-utan and rhesus macaque) in order to infer the reduction in allelic diversity that exists in humans as a consequence of purifying selection on linked sites over evolutionary timeframes^48^. As the effects of background selection on large genomic regions can mimic those of positive selection^46^, it is possible that the B-statistic might amalgamate both, though the rather large diversity reduction that it infers for the human genome as a whole suggests any bias due to positive selection is likely to be minor^44^. Finally, XP-EEH is a haplotype-based statistic which compares two population samples, and thus its power is increased for alleles that have suffered differential selective pressures since those populations diverged^105^. Though methodologically different, LA has a similar rationale by comparing human and Neanderthal genomes^107^, in order to infer the probability of each human haplotype to have been the result of an admixture event with Neanderthals.

For this work, CLR, CMS, B-statistic and LA were retrieved directly from their published references, and lifted over to GRC37 genomic coordinates if required using the ENSEMBL LiftOver tool^110,111^. As the available genome-wide measures of iHS and XP-EEH were based on HapMap3 data^112^, both statistics were re-calculated with the HAPBIN^113^ software directly on the EUR superpopulation of the 1KGPp3 dataset, with the AFR superpopulation used as the second population for XP-EEH. Taking advantage of the fine-scale genomic resolution of these statistics (between 1-10 kb), all SNP positions present in CLOZUK + PGC were assigned a value for each measure, either directly (if the position existed in the lifted-over data) or by linear interpolation. To simplify the interpretation of our results, all measures were transformed before further analyses to a common scale, in which larger values indicate stronger effect of selection or increased probability of introgression.

Heritability enrichment of these statistics was tested by the LD-Score partitioned heritability procedure. We derived binary annotations from the natural selection metrics by dichotomising at extreme cut-offs defined by the top 2%, 1% and 0.5% of the values of each metric in the full set of SNPs. This approach is widely used in evolutionary genomics, due to the difficulty of setting specific thresholds to define regions under selection^25,51^. Consistent with the previously described LDSR partitioned heritability protocol, enrichment was estimated for all binary annotations after controlling for 3 main categories of our set-based analysis (“non-genic”, “genic” and “genic LoF-intolerant”) and the 53 “baseline” categories of Finucane et al. 2015^22^. We note that the LD-Score software allows use of an extension of the partitioned heritability framework to test the metrics as fully quantitative annotations. Results of this analysis replicated those reported here (*data not shown*), though it did not constitute our main approach as methodological details on this feature of LDSR are yet to be published.

## ACKNOWLEDGEMENTS

General

This project has received funding from the European Union’s Seventh Framework Programme for research, technological development and demonstration under grant agreement n° 279227 (CRESTAR Consortium; http://www.crestar-project.eu/). The work at Cardiff University was funded by Medical Research Council (MRC) Centre (MR/L010305/1), Program Grant (G0800509) and the European Community’s Seventh Framework Programme HEALTH-F2-2010-241909 (Project EU-GEI). U.D. received funding from the German Research Foundation (DFG; FOR 2107 grant no. 1151/5-1).

Case data

We thank the participants and clinicians who took part in the CardiffCOGS study. For the CLOZUK2 sample we thank Leyden Delta for supporting the sample collection, anonymisation and data preparation (particularly Marinka Helthuis, John Jansen, Karel Jollie and Anouschka Colson), Magna Laboratories, UK (Andy Walker) and, for CLOZUK1, Novartis and The Doctor’s Laboratory staff for their guidance and cooperation. We acknowledge Lesley Bates, Catherine Bresner and Lucinda Hopkins, at Cardiff University, for laboratory sample management. We acknowledge Wayne Lawrence and Mark Einon, at Cardiff University, for support with the use and setup of computational infrastructures.

Control data

A full list of the investigators who contributed to the generation of the Wellcome Trust Case Control Consortium (WTCCC) data is available from www.wtccc.org.uk. Funding for the project was provided by the Wellcome Trust (WT) under award 076113. The UK10 K project was funded by the Wellcome Trust award WT091310. Venous blood collection for the 1958 Birth Cohort (NCDS) was funded by the UK’s Medical Research Council (MRC) grant G0000934, peripheral blood lymphocyte preparation by Juvenile Diabetes Research Foundation (JDRF) and WT and the cell-line production, DNA extraction and processing by WT grant 06854/Z/02/Z. Genotyping was supported by WT (083270) and the European Union (EU; ENGAGE: HEALTH-F4-2007-201413). The UK Blood Services Common Controls (UKBS-CC collection) was funded by WT (076113/C/04/Z) and by the National Institute for Health Research (NIHR) programme grant to NHS Blood and Transplant authority (NHSBT; RP-PG-0310-1002). NHSBT also made possible the recruitment of the Cardiff Controls, from participants who provided informed consent. Generation Scotland (GS) received core funding from the Chief Scientist Office of the Scottish Government Health Directorates CZD/16/6 and the Scottish Funding Council HR03006. Genotyping of the GS:SFHS samples was carried out by the Genetics Core Laboratory at the WT Clinical Research Facility, Edinburgh, Scotland and was funded by the MRC. The Type 1 Diabetes Genetics Consortium (T1DGC; EGA dataset EGAS00000000038) is a collaborative clinical study sponsored by the National Institute of Diabetes and Digestive and Kidney Diseases (NIDDK), National Institute of Allergy and Infectious Diseases (NIAID), National Human Genome Research Institute (NHGRI), National Institute of Child Health and Human Development (NICHD), and JDRF. The People of the British Isles project (POBI; http://www.peopleofthebritishisles.org) is supported by WT (088262/Z/09/Z). TwinsUK is funded by WT, MRC, EU, NIHR-funded BioResource, Clinical Research Facility and Biomedical Research Centre based at Guy’s and St Thomas’ NHS Foundation Trust in partnership with King’s College London. Funding for the QIMR samples was provided by the Australian National Health and Medical Research Council (241944, 339462, 389875, 389891, 389892, 389927, 389938, 442915, 442981, 496675, 496739, 552485, 552498, 613602, 613608, 613674, 619667), the Australian Research Council (FT0991360, FT0991022), the FP-5 GenomEUtwin Project (QLG2-CT-2002-01254) and the US National Institutes of Health (NIH; AA07535, AA10248, AA13320, AA13321, AA13326, AA14041, MH66206, DA12854, DA019951), and the Center for Inherited Disease Research (Baltimore, MD, USA). TEDS is supported by a program grant from the MRC (G0901245-G0500079), with additional support from the NIH (HD044454; HD059215). In the GERAD1 Consortium, Cardiff University was supported by WT, MRC, Alzheimer’s Research UK (ARUK) and the Welsh Government. Kings College London acknowledges support from the MRC. The University of Belfast acknowledges support from ARUK, Alzheimer’s Society, Ulster Garden Villages, N.Ireland R&D Office and the Royal College of Physicians/Dunhill Medical Trust. Washington University was funded by NIH grants, Barnes Jewish Foundation and the Charles and Joanne Knight Alzheimer’s Research Initiative. The Bonn group was supported by the German Federal Ministry of Education and Research (BMBF), Competence Network Dementia and Competence Network Degenerative Dementia, and by the Alfried Krupp von Bohlen und Halbach-Stiftung.

